# Antiviral roles of interferon regulatory factor (IRF)-1, 3 and 7 against human coronavirus infection

**DOI:** 10.1101/2022.03.24.485591

**Authors:** Danyang Xu, Joseph K. Sampson Duncan, Maria Licursi, Jin Gohda, Yasushi Kawaguchi, Kensuke Hirasawa

**Affiliations:** Division of BioMedical Sciences, Faculty of Medicine, Memorial University of Newfoundland, St. John’s, NL, Canada, A1B 3V6; Research Center for Asian Infectious diseases, The Institute of Medical Science, The University of Tokyo; Division of Molecular Virology, Department of Microbiology and Immunology, The Institute of Medical Science, The University of Tokyo, Tokyo, Japan; Department of Infectious Disease Control, International Research Center for Infectious Diseases, The Institute of Medical Science, The University of Tokyo

## Abstract

Interferon regulatory factors (IRFs) are key elements of antiviral innate responses that regulate transcription of interferons (IFNs) and IFN-stimulated genes (ISGs). As many human coronaviruses are known to be sensitive to IFN, antiviral roles of IRFs are yet to be fully understood. TypeI or II IFN treatment protected MRC5 cells from infection of human coronavirus 229E, but not human coronavirus OC43. Infection of 229E or OC43 efficiently upregulated ISGs, indicating that antiviral transcription is not suppressed during their infection. Antiviral IRFs, IRF1, IRF3 and IRF7, were activated in cells infected with 229E, OC43 or severe acute respiratory syndrome-associated coronavirus 2 (SARS-CoV-2). RNAi knockdown and overexpression of the IRFs demonstrated that IRF1 and IRF3 have antiviral property against OC43 while only IRF3 and IRF7 are effective to restrict 229E infection. Our study demonstrates that IRF3 plays critical roles against infection of human coronavirus 229E and OC43, which may be an anti-human coronavirus therapeutic target.

## Introduction

Human coronaviruses are enveloped single-stranded RNA viruses with positive-sense genomes that commonly cause respiratory tract infection in humans (1,2). They are comprised of 4 genera: alphacoronavirus, betacoronavirus, gammacoronavirus, and deltacoronavirus. Certain betacoronaviruses are known to cause lethal infection in humans, including middle east respiratory syndrome (MERS), severe acute respiratory syndrome-associated coronavirus (SARS-CoV) and SARS-CoV-2. MERS infection was first found in 2012; since then, 2249 infections and 858 deaths in 27 countries have been reported (3). SARS-CoV caused 8237 infections and 775 deaths in more than 30 countries in 2002 (4). SARS-CoV-2 was identified in 2019 and has caused the current COVID-19 pandemic. As of today (March 1, 2022), 437 million cases and 5.9 million deaths have been reported worldwide (5). Other human coronaviruses, such as OC43, 229E, NL63 and HKUI1, infect the upper respiratory tract and cause common seasonal cold symptoms (6). OC43 and HKUI1 belong to betacoronaviruses while 229E and NL63 are alphacoronaviruses (7,8). As the sequence of non-structural proteins are well-conserved among human coronaviruses, they share very similar replication cycles (9,10).

Cells sense viral products intracellularly and extracellularly using different pattern recognition receptors (PRRs) such as toll like receptors, RIG-I-like receptors and melanoma differentiation-associated gene 5 (MDA-5) (11). The recognition of viral products results in the nuclear translocation of IFN regulatory factor 3 (IRF3), IFN regulatory factor 7 (IRF7) and nuclear factor-κB (NF-κB), which activate the transcription of interferons (IFNs) (12,13). IFNs, which have three classes, type I (IFN-α/β), type II (IFN-γ) and type III (IFN-λ) IFNs, play essential roles in antiviral innate immune response (14,15). Secreted IFNs bind to IFN receptors in an autocrine or paracrine manner and activate the Janus kinase (JAK)-signal transducer and activator of transcription (STAT) (16,17). Phosphorylated STAT proteins directly bind to the promoter regions of IFN-stimulated genes (ISGs) to activate their transcription along with other transcriptional regulators such as IRF1 and IRF9 (18,19). These ISGs function as antiviral factors and immune regulators.

Human coronaviruses are generally sensitive to antiviral actions of IFNs albeit with some differences in their sensitivity. Both SARS-CoV and MERS are sensitive to IFN when cells are treated at high concentrations (20–23). Between the two viruses, IFNs are more effective in inhibiting the replication of MERS than SARS-CoV (23). Moreover, SARS-CoV-2 is much more sensitive to type I IFN than SARS-CoV (24,25). As for other human coronaviruses, IFNs suppress OC43 infection in a cell type dependent manner (26), while 229E is sensitive to IFN treatment *in vitro* (27,28). These reports suggest that human coronaviruses are generally sensitive to IFNs but each virus has different levels of sensitivity.

In clinical settings, IFN-β treatment significantly reduced the mortality of SARS-CoV-2 infected patients when administrated at an early stage of infection (29). Similarly, treatment of pegylated IFN-α significantly reduced viral replication of SARS-CoV in macaques (30). In STAT1 -/-mice, SARS-CoV induced a prolonged infection with higher viral loads in the lung, suggesting that the JAK/STAT pathway downstream of IFN receptors is essential for clearing SARS-CoV *in vivo* (31). However, SARS-CoV infection was not exacerbated in IFN receptor -/-mice but instead mouse survival was improved due to reduced immune cell infiltration in the lung, indicating immunopathogenic roles of IFNs in SARS-CoV infection (32).

While the antiviral efficacy of IFNs against human coronavirus is clear, SARS-CoV-2 infected patients only showed a low production of type I and III IFN and a moderate ISG response (33). Similarly, type I IFN response was delayed in mice infected with SARS-CoV-2, allowing viral replication, lung immunopathogenesis and lethal pneumonia (34). These reports suggest that IFN-mediated antiviral innate response is dysregulated in SARS-CoV-2 infection *in vivo*. This is most likely due to the presence of SARS-CoV-2 proteins that supress the production of IFNs and IGSs (35). In summary, IFNs have both antiviral and immunopathogenic roles in human coronavirus infection. Moreover, IFN antiviral responses are targets of immune evasion mechanisms by human coronaviruses.

Among the family of IRFs, IRF1, IRF3 and IRF7 are transcriptional regulators of IFNs and ISGs (36,37). IRF3 can be important for innate immune responses against SARS-CoV-2, as blocking phosphorylation and translocation of IRF3 promotes its replication (38). Accumulating evidence suggests that human coronaviruses can interfere with the activity of IRF3. SARS-CoV-2 PLpro and 3CLpro, viral proteins responsible for cleaving viral polyproteins, also degrade IRF3 (39,40). Other studies demonstrated that SARS-CoV-2 7a reduces IRF3 phosphorylation by downregulating TANK Binding Kinase 1 (TBK1) expression levels (41,42). Similarity, SARS-CoV 8b and 8ab induce IRF3 degradation on a ubiquitin dependent manner (43), while MERS M disrupts interaction of TNF Receptor Associated Factor 3 (TRAF3) and TBK1, leading to reduced IRF3 activation (43). These studies clearly suggest that human coronavirus proteins target IRF3 to promote their replication. In contrast to IRF3, antiviral roles of IRF1 and IRF7 against human coronavirus infection are less understood. In animal coronaviruses, IRF1 was shown to have antiviral properties against mouse hepatitis virus (MHV) (44). M protein of porcine epidemic diarrhea virus (PEDV) interacts with IRF7 and inhibits its antiviral functions (45,46). Thus, it is possible that IRF1 and IRF7 also have antiviral effects in human coronavirus infection.

Although it is suggested that the antiviral IRFs are important for host antiviral responses against human coronavirus infection, there is no direct functional evidence reported. In this study, to clarify antiviral functions of IRF1, IRF3 and IRF7 during human coronavirus infection, we conducted loss- and gain-of-function experiments of IRF1, IRF3 and IRF7. To fight against current and future pandemics, it is important to grasp more knowledge about antiviral responses mediated by the IRFs during human coronavirus infection.

## Materials and methods

### Cells, viruses, and reagents

Human lung fibroblast cell line MRC5, human lung cancer cell line H1299, human lung cancer cell line Calu-3, mouse fibroblast cell line L929, human coronaviruses HCoV-OC43 and HCoV-229E were obtained from the American Type Culture Collection (ATCC; Manassas, VA, USA). Vesicular stomatitis virus (VSV, Indiana strain) was provided by Dr. John C. Bell (Centre for Innovative Cancer Therapeutics, Ottawa Hospital Research Institute, Ottawa, Canada). VSV was amplified and titrated by plaque assay using L929 cells as described previously (47). Recombinant human IFN-α A, human IFN-γ and IFN-λ 1 were obtained from Bio-Rad, BD Pharmingen and R&D Systems respectively. Antibodies used in this study include: IRF3, phospho-IRF3, IRF7, phsopho-STAT1 (Cell signalling technology), IRF1 (BD Transduction Laboratories), GAPDH (Santa Cruze Biotechnology), 229E N protein (Ingenasa), OC43 N protein (Milipore), SARS-CoV-2 N protein (Sino Biological). Negative control siRNA, IRF1 siRNA (s7501), IRF3 siRNAs (s7509) and IRF7 siRNA (s223948) were purchased from Life Technologies. IRF1-pINCY plasmid (Open biosystems) and IRF7 -ORF vector (Applied Biological Materials) was subcloned into pcDNA3 plasmid (Addgene). pcDNA3-IRF3 was purchased from Addgene.

### Cell culture

All cells were cultured in high-glucose Dulbecco’s modified Eagle’s medium (Corning, MA) with 10% fetal bovine serum (HyClone, Cytiva), 1 mM sodium pyruvate and antibiotic-antimycotic (Life Technologies).

### Virus infection

Cells with 90% confluency were infected with human coronavirus 229E or OV43 at MOI of 0.01. The diluted stock viruses were absorbed for 2 hours at 33 °C, and then removed and replaced with DMEM with 2% FBS. Infected cells were incubated at 33 °C with 5% CO_2_ for up to six days. For IFN treatment, MRC5 cells were treated with IFN-α (250 and 500 U/ml), IFN-γ (50 and 100 U/ml) or left untreated for 18 hours and then challenged with 229E or OC43. For siRNA knockdown, cells were transfected with 5 pmol siRNA oligoes using Lipofectamine RNAiMAX Transfection Reagent (Life Technologies) and 24 hours later challenged with or without 229E or OC43. For overexpression of IRFs, cells in 24-well plates (4×10^4^ cells/well) were transfected with control pcDNA3, pcDNA-IRF1, pcDNA-IRF3 or pcDNA-IRF7 using Lipofectamine 3000 Transfection Reagent (Life Technologies) and 24 hours later challenged with or without 229E or OC43. VSV absorption and infection (MOI of 0.0001) was conducted at 37 °C with 5% CO_2_. The amount of progeny viruses in the culture supernatant was measured by TCID_50_ (50% tissue culture infective dose) assay for 229E and OC43 (48) and plaque assay for VSV (47). The SARS-CoV-2 isolate (UT-NCGM02/Human/2020/Tokyo) (49) was propagated in VeroE6-TMPRSS2 cells in DMEM containing 5% heat-inactivated FBS at 37 °C in 5 % CO2. Calu-3 cells were infected with SARS-CoV-2 at a MOI of 0.1 and absorbed for 30 min at 37°C. Then, the culture medium was replaced with fresh medium.

### Western blot analysis

Western blot analysis on cell lysates was conducted and evaluated using antibodies listed above as previously described (47).

### Quantitative RT-PCR

Quantitative RT-PCR (RT-qPCR) was performed in triplicate using the previously described validation strategies (50). The primer sequences are shown in Supplementary Table S1.

### Statistical analysis

Two-way ANOVA followed by Sidak’s post hoc test was performed using GraphPad Prism 6.0 software.

## Results

### IFN treatment delays 229E infection but not OC43 infection

To investigate the antiviral effects of IFNs on human coronavirus infection, we first tested if human lung fibroblast cells, MRC5, are sensitive to different types of IFNs (Figure 1A). When MRC5 cells were treated with IFN-α, IFN-γ or IFN-λ for 30 mins, we observed STAT1 phosphorylation in cells treated with IFN-α or IFN-γ, but not in those treated with IFN-λ. This indicates that MRC5 cells are sensitive to type I and type II IFN, but not to type III IFN. Therefore, we focused on type I and II IFN for the following experiments.

**Figure 1:**
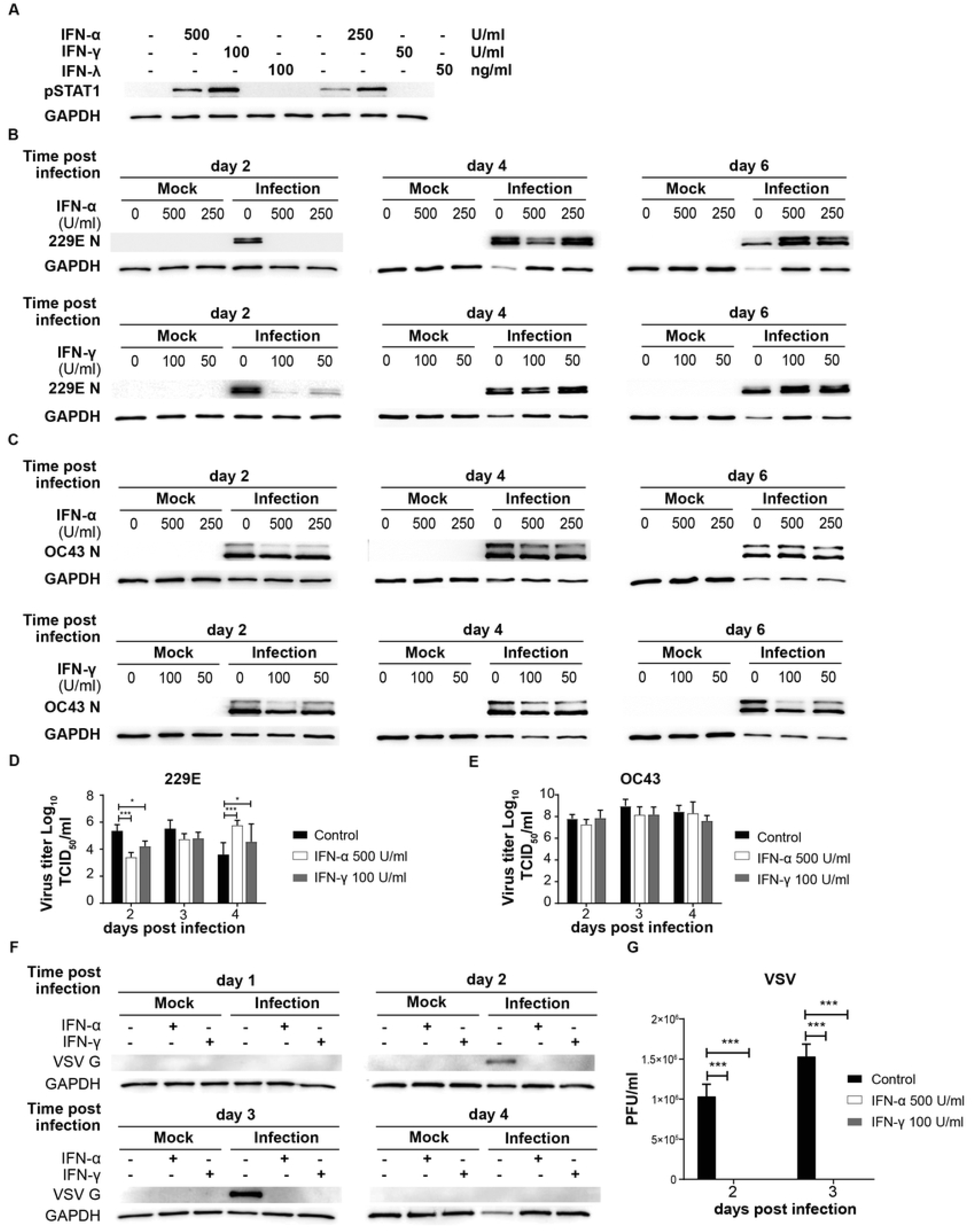
IFN treatment delays 229E but not OC43 infection. (A) MRC5 cell were left untreated or treated with IFN-α (500 and 250 U/ml), IFN-γ (100 and 50 U/ml) or IFN-λ (100 and 50 ng/ml). STAT1 activation was determined by western blot analysis using anti phospho-STAT1 and GAPDH antibodies. (B and C) MRC5 cells were left untreated or treated with IFN-α or IFN-γ, and then challenged with 229E (B) or OC43 (C) infection at MOI of 0.01. Western blot analysis of viral protein was conducted using anti 229E N protein (B), OC43 N protein (C) and GAPDH antibodies. (D and E) TCID_50_ assay was performed to measure the progeny virus of 229E or OC43 infected MRC5 cells left untreated or treated with IFNs. (F and G) MRC5 cells were left untreated or treated with IFN-α (500U/ml) or IFN-γ (100U/ml), and then challenged with VSV at MOI of 0.0001. VSV infection was determined by (F) western blot analysis using anti VSV G and GAPDH antibodies and (G) plaque assay using L929 cells. The amount of progeny virus is shown as plaque forming units (PFU)/ml of samples nontreated or treated with IFN. *p<0.05, **p<0.01, ***p<0.001, ****p<0.0001, Two-way ANOVA.

MRC5 cells were left untreated or treated with IFN-α (250 and 500 U/ml) or IFN-γ (50 and 100 IU/ml) for 18 hours and then challenged with 229E (Figure 1B) or OC43 (Figure 1C) at MOI of 0.01. Cell lysates were harvested at 2, 4 and 6 days after infection for western blot analysis of viral nucleocapsid proteins (OC43 N and 229E N) and GAPDH. At 2 days after 229E infection, viral protein was detected in cells without IFN-α treatment, but not in cells treated with IFN-α (Figure 1B). At 4 days after infection, less viral protein was detected with a higher IFN-α concentration (500 IU/ml). Similarly, IFN-γ treatment inhibited 229E infection at 2 days, but not at 4 and 6 days after infection. In contrast, OC43 infection was not significantly affected by IFN-α or IFN-γ, although some minor reductions in viral protein levels were observed in cells treated with IFNs at 2 days after infection (Figure 1C). To further confirm the effect of IFNs on virus production, we conducted a progeny virus assay (Figure 1D and E). The amount of progeny 229E was significantly lower in cells treated with IFN-α or IFN-γ at 2 days after infection (Figure 1D). On the other hand, IFN treatment did not reduce the progeny virus production of OC43 (Figure 1E). To confirm the efficacy of IFN to induce sufficient antiviral responses, we conducted a positive control experiment where MRC5 cells were treated with the same amount of IFN-α or IFN-γ, and then challenged with an IFN-sensitive virus, vesicular stomatitis virus (VSV). The IFN treatment completely inhibited VSV protein synthesis (Figure 1F) and progeny virus production (Figure 1G), indicating the concentration of IFN used in our system is sufficient to inhibit replication of an IFN sensitive virus.

Taken together, these results suggest that both type I and II IFN delay 229E infection in MRC5 cells but they are not effective in protecting against OC43 infection.

### 229E and OC43 infection activate transcription of IFN-stimulated genes

Our next question was whether human coronaviruses activate antiviral innate responses in infected cells. To test this, we assessed the transcriptional activation of ISGs at 2 and 4 days after human coronavirus infection (Figure 2). Western blot analysis was first conducted to confirm infection of 229E (Figure 2A) and OC43 (Figure 2B). Then, the expression of the following IFN-inducible genes was examined during 229E (Figure 2C) or OC43 (Figure 2D) infection: guanylate binding protein 2 (GBP2) (51), interferon induced protein 44 (IFI44) (52), interferon induced protein with tetratricopeptide repeats 2 (IFIT2) (53), microtubule-associated protein 2 (MAP2) (54), retinoic acid-inducible gene I (RIG-I) (55) and STAT2 (56). 229E infection did not induce GBP2, but significantly increased the expression of all other genes at 4 days after infection. In contrast, OC43 infection induced all IFN-inducible genes tested at 2 days after infection, and the expression levels were significantly higher than control at 4 days except for GBP2 and STAT2. These results indicate that human coronavirus infection efficiently induces host antiviral transcriptional responses.

**Figure 2:**
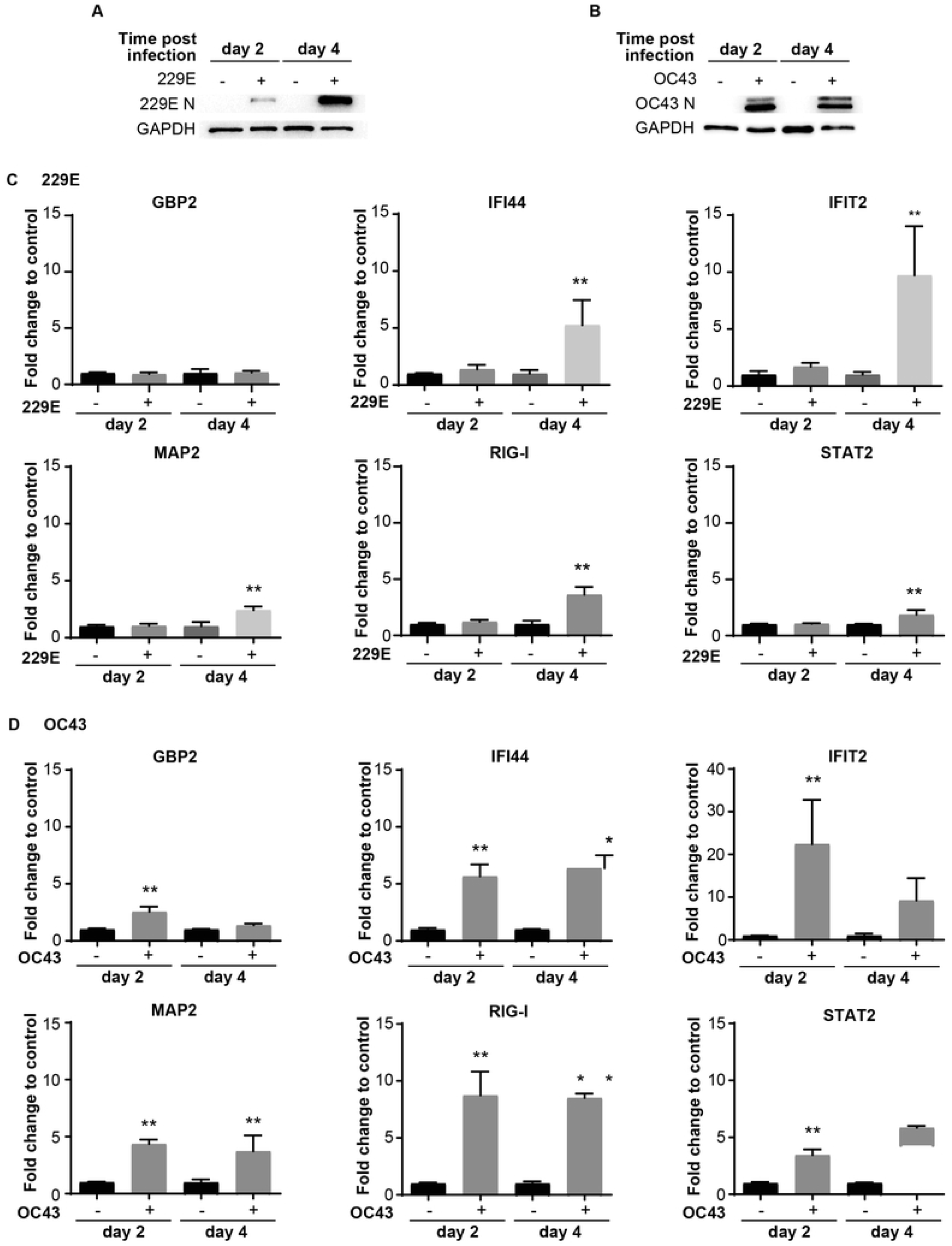
Human coronavirus infection activates transcription of ISGs. MRC5 cells were left uninfected or infected with 229E (A and C) or OC43 (B and D) at MOI of 0.01. Infection of 229E (A) and OC43 (B) was confirmed by western blot analysis using anti 229E N protein, OC43 N protein and GAPDH antibodies. The mRNA levels of ISGs (GBP2, IFI44, IFIT2, MAP2, RIG-I and STAT2) were determined by RT-qPCR in MRC5 cells infected with 229E (C) or OC43 (D). The relative quantification (RQ) indicates the fold change of the expression level for infected samples towards that of the non-infected controls at the same time point post infection. The transcriptional level for each gene was calculated by normalizing to GAPDH expression level and then normalized by the corresponding control. **p<0.01, Two-way ANOVA.

### IRF1, IRF3 and IRF7 are activated during human coronavirus infection

As IRF1, IRF3 and IRF7 are the key transcriptional regulators of IFNs and ISGs, we sought to determine their activation status during human coronavirus infection. Accordingly, a western blot analysis was conducted to assess the expression of IRF1 and IRF7, and phosphorylation of IRF3 (an active form of IRF3) in MRC5 cells infected with 229E or OC43. After 229E infection, viral proteins were detected from day 2 and reached a peak at day 3 and 4 (Figure 3A). The expression of IRF1 and IRF7 was increased at 3 and 4 days after infection compared to uninfected controls. Similarly, phosphorylated IRF3 was increased at the same time points. After OC43 infection, OC43 nucleoprotein was detected on day 1, which peaked at 3 and 4 days after infection (Figure 3B). In these infected cells, IRF1 expression increased at day 2 and 3. We observed an upper shift in IRF1 bands, which may be caused by posttranslational modifications of IRF1. IRF3 and IRF7 were also activated from 2 to 5 days after OC43infection.

**Figure 3:**
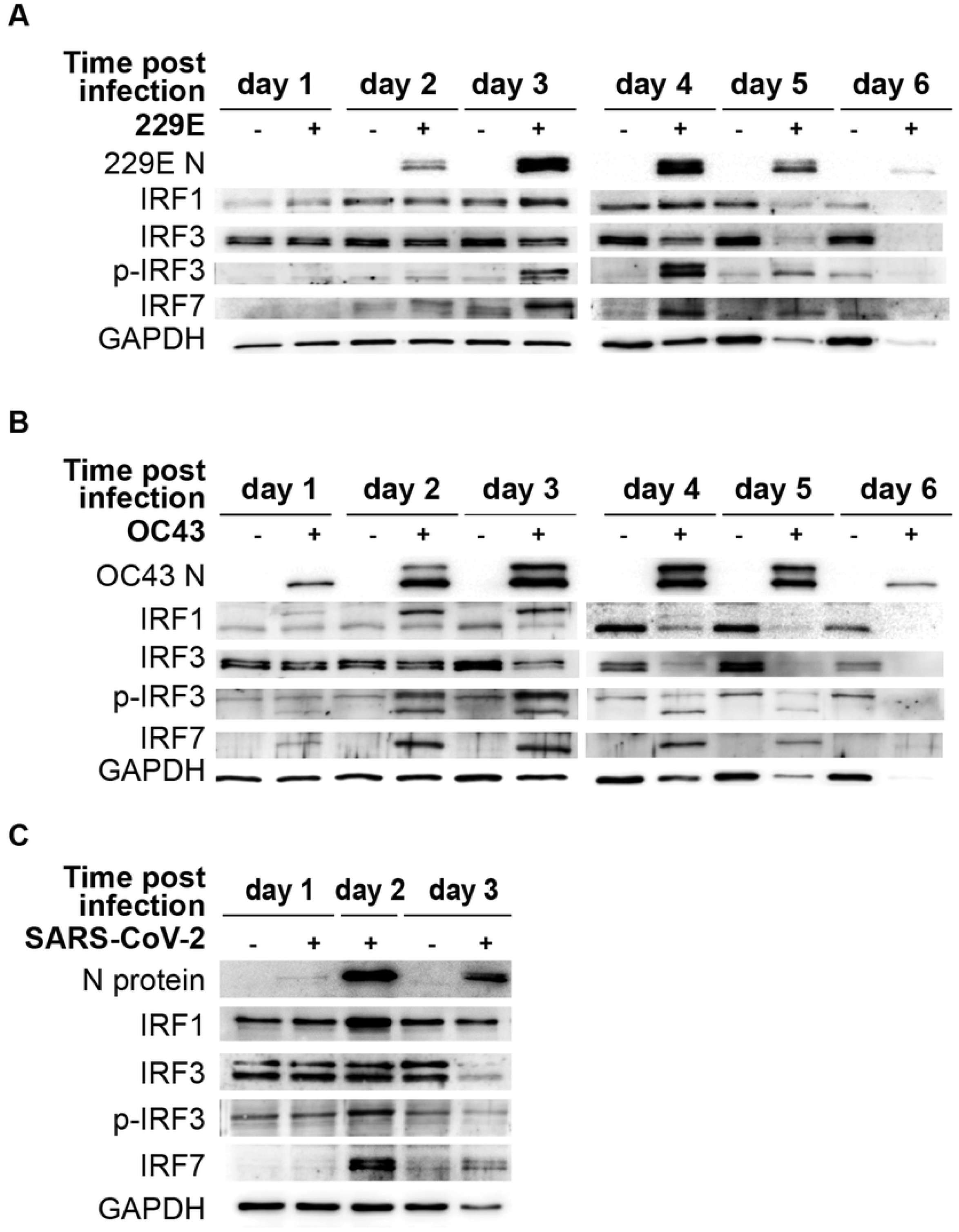
Human coronavirus infection activates IRF1, IRF3 and IRF7. MRC5 cells were left uninfected or infected with 229E (A) or OC43 (B) at MOI of 0.01. (C) Calu-3 cells were left uninfected or infected with SARS-CoV-2 at MOI of 0.001. The activation of IRF1, 3 and 7 was determined by western blot analysis using antibodies against 229E N protein, OC43 N protein, SARS-CoV-2 N protein, IRF1, IRF3, IRF7, phosphorylated (p-IRF3) and GAPDH.

To determine the effect of SARS-CoV-2 infection on IRFs, Calu-3 cells were infected with SARS-CoV-2. We detected viral protein after 1 day, which peaked at 2 days after infection (Figure 3C). In these cells, there was an increase in IRF1 and IRF7 expression and IRF3 phosphorylation at 2 days after infection. Thus, SARS-CoV-2 infection also activates antiviral IRFs.

These results demonstrate that IRF1, IRF3 and IRF7 are activated during human coronavirus infection.

### IRF1, 3 and 7 have antiviral roles against human coronavirus infection

To investigate the functional roles of IRF1, IRF3 and IRF7 during human coronavirus infection, we conducted a loss-of-function analysis using siRNA knockdown. MRC5 cells were transfected with either control siRNA oligos or those against IRF1, 3 or 7 for 24 hours. The knockdown of IRF1 and IRF3 was confirmed with western blot analysis showing lower expression levels in cells treated with their corresponding siRNA oligoes (Figure 4A). As IRF7 expression is undetectable by western blot in non-infected cells, IRF7 knockdown was confirmed by qPCR analysis (Figure 4B). When these cells were challenged with 229E or OC43, an RT-q-PCR analysis revealed that IRF1 knockdown promotes 229E infection at 3 days after infection and OC43 infection at 2 and 3 days after infection (Figure 4C). IRF3 knockdown also increased the expression of viral RNA at 2 and 3 days after 229E infection and 2 days after OC43 infection. In addition, IRF7 knockdown resulted in an increase in 229E infection at 3 days after infection and OC43 infection at 2 and 3 days after infection. These results were further confirmed by a western blot analysis (Figure 4 D). 229E viral protein synthesis was increased in cells treated with siRNA oligoes against IRF3 or IRF7 compared to siRNA controls at 2 days after infection. For OC43 infection, knockdown of IRF1, IRF3 or IRF7 promoted viral protein synthesis compared with siRNA controls on both 2 and 3 days after infection. Altogether, these loss-of-function experiments indicate the antiviral roles of IRF1, IRF3 and IRF3 against coronavirus infection.

**Figure 4:**
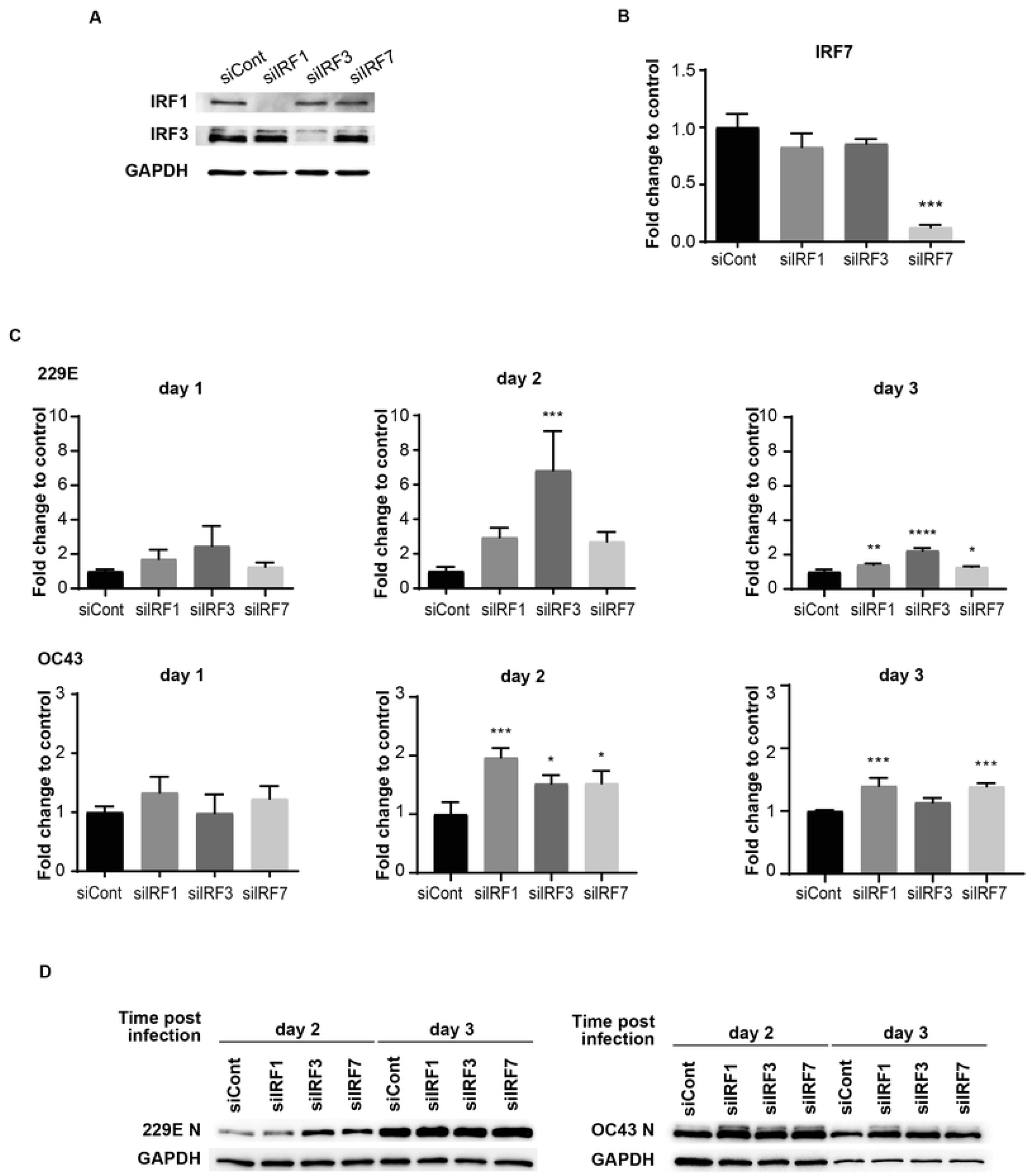
Knockdown of IRFs promotes human coronavirus infection. MRC5 cells were transfected with control siRNA (siCont), IRF1 siRNA, IRF3 siRNA or IRF7 siRNA oligoes (5 pmol) using Lipofectamine RNAiMAX transfection reagent. The knockdown of IRFs expression was confirmed by western blot analysis for IRF1 and IRF3 (A) and RT-qPCR for IRF7 (B). MRC5 cells were then infected with 229E or OC43 at MOI of 0.01. (C) The amounts of viral RNA were measured by RT-qPCR. (D) The amounts of viral protein were determined by western blot analysis using antibodies against 229E N protein, OC43 N protein and GAPDH. For RT-qPCR analysis, the transcription level for each gene was first normalized to GAPDH expression level. The relative quantification (RQ) indicates the fold change of the expression level for IRFs siRNA transfected samples to that of the control siRNA transfected samples. *p<0.05, **p<0.01, ***p<0.001, ****p<0.0001, Two-way ANOVA.

To further confirm the antiviral roles of IRF1, IRF3 and IRF7, we conducted their gain-of-function experiments. H1299 cells were transfected with control pcDNA3, IRF1-pcDNA3, IRF3-pcDNA3 or IRF7-pcDNA3 for 24 hours and then challenged with 229E or OC43 (Figure 5). IRF1 overexpression effectively inhibited replication of OC43 as the generation of viral proteins and progeny viruses were lower in cells transfected with IRF1-pcDNA than those transfected with control pcDNA3 (Figure 5A). We observed a slight reduction of 229E N protein expression in IRF1 overexpressed cells at 1 day after infection, but there was no significant difference in the amount of progeny virus (Figure 5D). Moreover, 229E and OC43 generated less viral proteins and progeny viruses in the cells transfected with IRF3-pcDNA3 (Figure 5B and E), suggesting that IRF3 introduction promoted antiviral activities against both 229E and OC43. Finally, the introduction of IRF7 was effective to reduce 229E infection but not OC43 infection as shown in western blotting and progeny virus analysis (Figure 5C and F).

**Figure 5:**
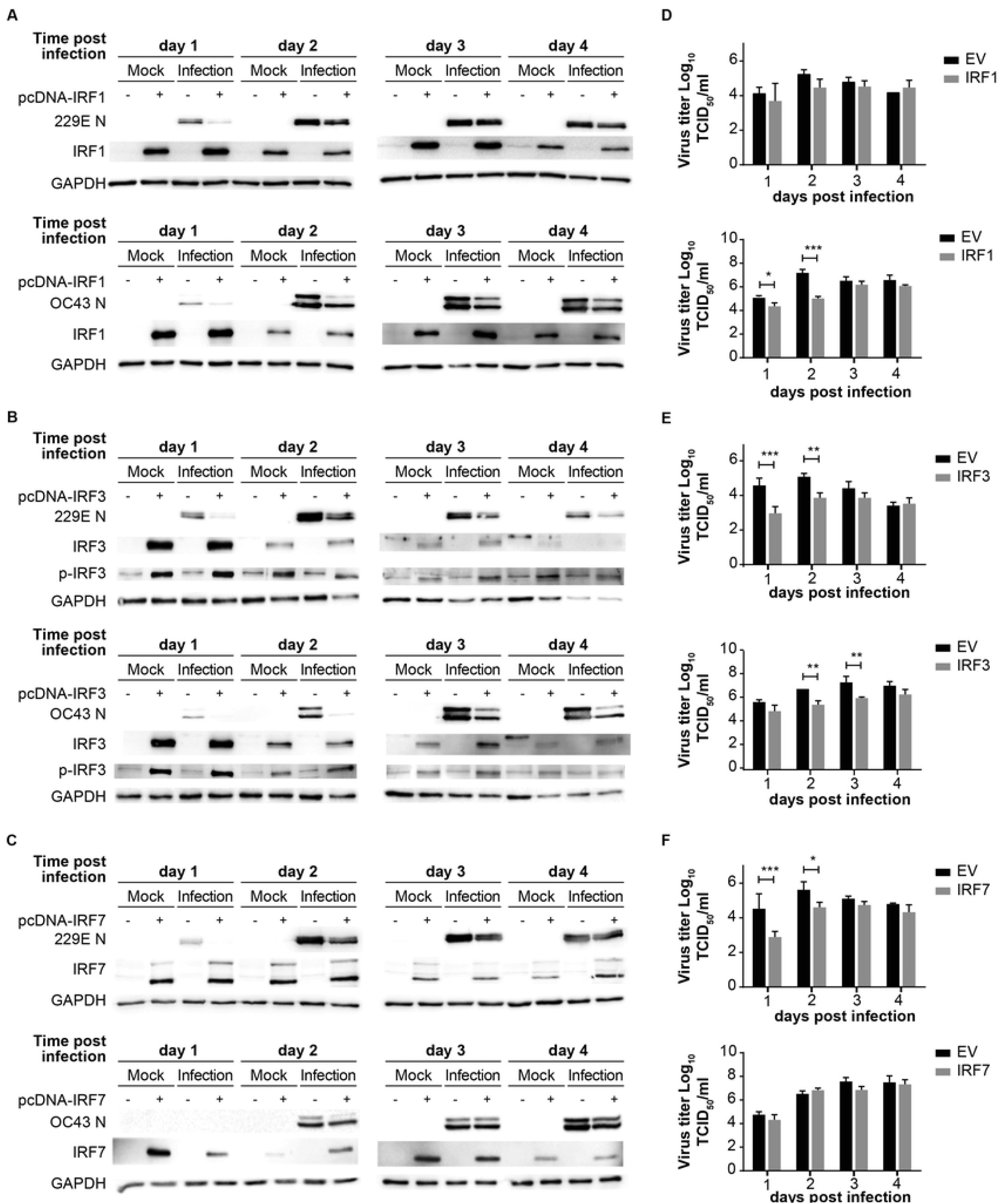
Overexpression of IRFs inhibits human coronavirus infection. H1299 cells were transfected with control pcDNA3 plasmid or the plasmid containing IRF1 (A), IRF3 (B) or IRF7 (C) and then infected with or without 229E or OC43 at MOI of 0.01. Virus infection was determined by western blot analysis using antibodies against 229E N protein, OC43 N protein, IRF1, IRF3, IRF7, phosphorylated IRF3 (p-IRF3) and GAPDH. Amounts of progeny viruses were measured in the supernatant of cells transfected with the plasmid containing with IRF1 (D), IRF3 (E) or IRF7 (F) by TCID_50_ assay which were compared to in those transfected with empty plasmid. *p<0.05, **p<0.01, ***p<0.001, Two-way ANOVA

## Discussion

In this study, we demonstrated an antiviral role of the IFN-IRF axis against human coronavirus infection. We first found that human coronavirus 229E is moderately sensitive to type I and II IFN while OC43 is not (Figure 1). Infection of both viruses efficiently induced ISGs and activated IRF1, IRF3 and IRF7, suggesting that the antiviral innate response of infected cells is not fully suppressed during infection (Figure 2 and 3). Activation of IRFs was also observed during SARS-CoV-2 infection (Figure 3). The loss- and gain-of function experiments demonstrated that IRF1 and IRF3 have antiviral roles against OC43 infection while IRF3 and IRF7 were effective to suppress 229E infection (Figure 4 and 5).

We found that 229E is sensitive to type I and II IFN, in agreement with previous studies (21,28). Type I IFN has been shown to inhibit OC43 infection in A549 (lung cancer cells) but to promote it in NCI-H520 (lung cancer cells) or Huh 7.5 (hepatoma) (57). In our current study, IFN treatment did not have any effect on OC43 replication in MRC5 (normal fibroblast cells). This discrepancy could be due to the differences in cell types. Alternatively, it may be because of IFN concentrations used in this study, which is lower than those used in previous studies. The concentrations of IFN in the present study were based on our previous work on IFN-sensitive viruses (47,58). A comparison in the same experimental system showed that the same concentration of IFN completely shuts down the replication of IFN-sensitive VSV, but only partially suppresses 229E while not affecting OC43 (Figure 1F and G). Thus, we conclude that the IFN sensitivity of human coronavirus 229E and OC43 is not very high.

Certain viral proteins of human coronaviruses are known to degrade IRF3 (40,43). This was the case in our study where the expression of IRF3 was decreased during infection of 229E or OC43 (Figure 3). Nevertheless, we found that IRF3 was efficiently phosphorylated during viral infection (Figure 3). Furthermore, siRNA knockdown of IRF3 increased the susceptibility of host cells to 229E or OC43 infection (Figure 4). These results indicate that IRF3-mediated antiviral response is still active in cells infected with human coronaviruses, although viral evasion downregulates its expression. It was shown previously that BX795, which blocks phosphorylation and translocation of IRF3 (59), inhibits the induction of ISGs and promotes replication of SARS-CoV-2 (60). Considering that IRF3 overexpression inhibited 229E or OC43 infection (Figure 5), IRF3 may be a common antiviral effector against human coronavirus infection, which would make it an excellent antiviral therapeutic target. In contrast, the promotion of IRF1 showed antiviral activities in cells infected with OC43 while that of IRF7 expression reduced 229E infection (Figure 5). We are not certain if this indicates specific antiviral roles of IRFs against different human coronaviruses. To this end, it is essential to further expand our study to investigate antiviral functions of the IRFs in other human coronaviruses.

IRF1, IRF3 and IRF7 are critical transcriptional regulators of IFNs and ISGs (61–63). Alternately, IFNs are major transcriptional activators of the IRFs (37). Thus, the activities of IFNs and IRFs are closely related. Interestingly, we found that OC43 is sensitive to antiviral effects of IRF1 and IRF3 while insensitive to IFNs. This may be because of viral immune evasion of OC43 on the JAK/STAT pathway and the PRR signaling pathway, which is often observed during human coronavirus infection (64,65). If this is the case, activation of IRFs at the downstream transcriptional levels may be more effective than stimulating upstream antiviral innate immunity. Alternatively, there may be differences in the modes of innate immunity activated by IFN treatment and IRF modulation used in the study. IFN treatment induces strong but transient antiviral responses while modulation of IRFs using RNAi knockdown or plasmid overexpression induces constant, prolonged effects. Therefore, it is possible that an extended period of IFN treatment with different doses may be effective on OC43 infection.

In western blot analysis, we observed the IRF1 bands, which are higher than expected, at 2 and 3 days after OC43 infection (Figure 3B). Interestingly, this shift was not observed in cells infected with 229E or SARS-CoV-2 infected cells. We believe that the size increase of IRF1 may be induced by its posttranslational modifications. The phosphorylation or monoubiquitination promotes the transcriptional activities of IRF1, suggesting that the upper bands of IRF1 could be the active form (66). While this may be the reason why IRF1 showed antiviral activities against OC43 infection but not 229E infection (Figure 5A), it is yet to be clarified why they were observed only in cells infected with OC43.

We use MRC5 cells for most of our studies (Figure 1-4) while H1299 cells were used for the gain-of-function experiments for IRF3s (Figure 5). This is because we encounter technical problems to achieve sufficient expression levels of the IRFs by transfecting the plasmids without causing cell morbidity or affecting virus infection in MRC5 cells.

## Acknowledgements

This work was supported by grants from the Canadian Institutes for Health Research (CIHR) (KH), the Natural Sciences and Engineering Research Council of Canada (NSERC) (KH), the Japan Agency for Medical Research and Development (AMED) (Grant Number JP20wm0125002) (YK and JG) and Memorial University of Newfoundland (KH). DYX was supported by the Dean’s fellowship from Faculty of Medicine, Memorial University of Newfoundland. Special thanks to Dr. Yoshihiro Kawaoka (Division of Virology, Department of Microbiology and Immunology, Institute of Medical Science, University of Tokyo) for SARS-CoV-2 and Dr. John Bell (University of Ottawa, Ottawa, Canada) for VSV.

---

**Table S1.**
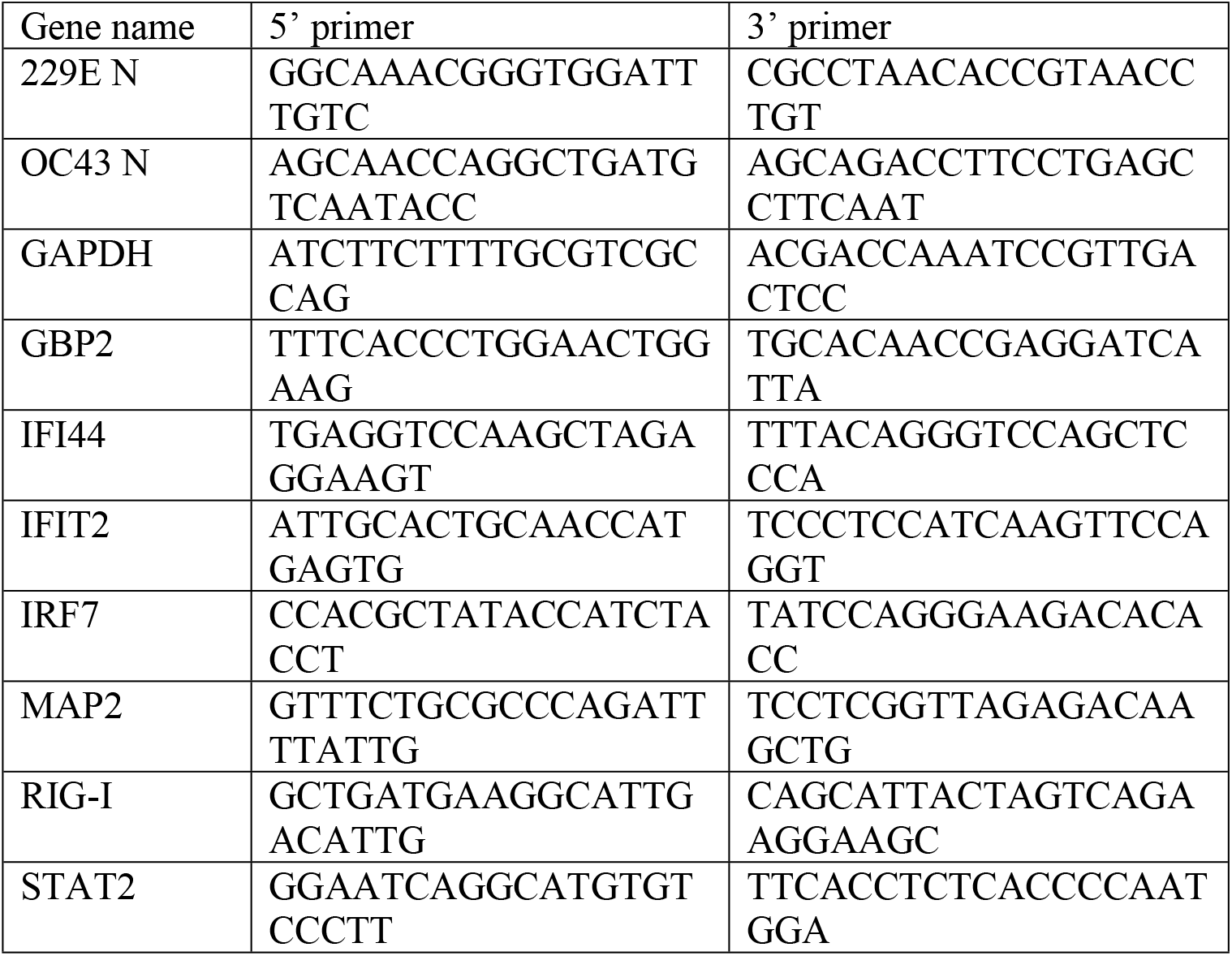
Sequence of qPCR primers

